# Folding the unfoldable 2: using AlphaFold and ESMFold to explore spurious proteins

**DOI:** 10.64898/2026.06.09.728211

**Authors:** Ailsa K. Orr, Alex Bateman

## Abstract

**Motivation:** Spurious protein sequences, resulting from gene prediction errors, theoretically should not yield folded structures. AlphaFold2 was previously shown to predict short spurious sequences with high pLDDT scores and was therefore unlikely to distinguish between real proteins and spurious proteins which are usually short. We evaluate whether newer structure prediction methods (ESMFold and AlphaFold3) similarly predict short sequences with high pLDDT or if they better discriminate between spurious and real proteins.

**Results:** All three structure prediction methods (ESMFold, AlphaFold2, and AlphaFold3) predict short spurious sequences from AntiFam with unexpectedly high pLDDT scores, however the discrimination between spurious and real proteins improves beyond 100 amino acids. By analysing sequences with disparate pTM and pLDDT scores, we identified two likely spurious shadow ORFs in Swiss-Prot and one potentially non-spurious AntiFam entry. Using the structure prediction scores, we developed a Gaussian Process Model and evaluated its performance on AlphaFold DB, identifying potential spurious proteins at scale. While limited on its own, this model can increase confidence in spurious protein identification when combined with other methods.

**Availability:** Structure predictions are available at https://doi.org/10.5281/zenodo.18390113. Model implementation and figure generation code are available at https://github.com/0rra/fold_unfold2.

## Introduction

Spurious proteins are the result of errors in protein-coding gene prediction. These erroneous entries arise from various sources, including non-coding RNA genes misidentified as protein coding (Tripp et al. 2011), genome contamination from repetitive elements (Breitwieser et al. 2019), or misidentification due to protein coding genes on the opposite strand (shadow ORFs), (Eberhardt et al. 2012; Trimble et al. 2012) or overlapping reading frames (Veloso et al. 2005).

As protein sequence databases continue to expand, the number of spurious proteins also increases. Only a small fraction of protein sequences is experimental verified or manually curated, the vast majority are automatically annotated. While an overall robust method, automatic gene prediction has a predicted error rate of 1-2% for bacterial genomes, which means potentially 2 to 5 million protein sequences in UniProtKB could be spurious.

AntiFam (Eberhardt et al. 2012) is a collection of profile hidden Markov models (HMMs) representing sequences believed to be spurious. It serves as a quality control tool for protein databases such as UniProtKB. With the release of version 8.0, AntiFam has grown to include 278 entries. AntiFam’s overall coverage of spurious proteins is likely to be low, but it is a useful training set for the development of more general methods to detect spurious proteins.

Previously AntiFam was used to evaluate AlphaFold2 (Jumper et al. 2021), and to investigate whether AlphaFold2 would assign globular structures to known spurious sequences (Monzon et al. 2022). That investigation revealed an important limitation: AlphaFold2 assigned high confidence scores (measured by predicted local distance difference test, pLDDT) to short AntiFam sequences, making them indistinguishable from real UniProtKB/Swiss-Prot proteins. This same effect was also found for randomly generated sequences. Additionally, one AntiFam entry (AntiFam: ANF00096, since removed) was identified as a false positive entry, containing real proteins rather than spurious ones.

Recent advances in protein structure prediction have introduced new methodologies: ESMFold (Lin et al. 2023), which employs a protein language model approach, removing the need for the time-consuming step of multiple sequence alignment creation, and AlphaFold3 (Abramson et al. 2024), which uses a generative diffusion approach distinct from AlphaFold2’s neural network-based architecture. These methodological differences provide an opportunity to investigate whether these newer tools can better distinguish between spurious and real protein sequences.

In this study, we aim to:

1. Evaluate whether AlphaFold3 and ESMFold predict high confidence structures for short spurious sequences, as previously observed with AlphaFold2
2. Identify any false positive AntiFam entries
3. Investigate whether structure prediction confidence scores can be used to train a machine learning model which distinguishes between real and spurious proteins

We assess performance using confidence metrics including pLDDT (which measures local structural quality on a scale of 0-100) and predicted TM-score (pTM, which indicates the predicted global structural similarity to the native structure).

## Methods

We used three types of input sequences for protein structure predictions: AntiFam sequences, randomly generated sequences, and UniProtKB/Swiss-Prot sequences (referred to as AntiFam, Random, and Swiss-Prot sequences, respectively).

AntiFam sequences were selected from AntiFam release 8.0, which includes 278 entries. Up to 3 sequences were selected from each AntiFam entry, with all sequences selected from smaller families of less than 3 sequences. Any gap characters within the sequences were removed, resulting in 763 AntiFam sequences.

To compare the prediction performance between spurious and real proteins, we randomly selected sequences from UniProtKB/Swiss-Prot (version release 2024_04). The protein sequences were divided into subsets of specific lengths between 10 to 200 (10, 16, 20, 30, 40, 50, 60, 70, 80, 90, 100, 120, 140, 160, 180, 200), and were then filtered to remove (i) fragment sequences and (ii) those with an average IUPred2A score above 0.5 to exclude disordered protein sequences. (Erdős and Dosztányi 2020; Mészáros et al. 2018). From this filtered set of non-fragmented and non-disordered proteins, five proteins of each length were randomly selected, totalling 80 Swiss-Prot sequences.

As an additional control we generated random sequences at defined lengths between 10 to 200, using the amino acid composition weighting from Swiss-Prot. Five sequences were generated per length, totalling 80 Random sequences.

We ran structure predictions within a NextFlow pipeline, capable of performing AlphaFold2 (ColabFold 2.3.6), AlphaFold3 (commit 7f6edd04) and ESMFold ((fair-esm) version 2.0.0) predictions. The NextFlow pipeline and code used for plots can be found on https://github.com/0rra/fold_unfold2. The prediction confidence scores, mean pLDDT, pTM, and PAE (available for AlphaFold2 and AlphaFold3), for each structure prediction were recorded.

We trained a Gaussian Process Classifier (GPC) model using the three features, length, mean pLDDT and mean pTM, to classify structures as spurious or real. Model training and testing metrics were calculated using Python 3.13, scikit-learn 1.61. For the training data set, we used one sequence per AntiFam entries, excluding ANF00264 (277 sequences), and 277 bacterial Swiss-Prot sequences. For the training set, we used only bacterial Swiss-Prot sequences with lengths less than or equal to 100 to improve the model’s ability to distinguish short sequences. To avoid data leakage of the Swiss-Prot training set to the Swiss-Prot testing set, we clustered the sequences with MMseqs2 (Mirdita et al. 2019), with coverage 90% and minimum identity 30%, and ensured that the sequence clusters used in the training dataset did not appear in the testing dataset.

Due to the limited number of AntiFams available, we generated AntiFam-like sequences to be used for model testing. These AntiFam-like sequences were made by translating protein-coding DNA of reference bacteria genomes in all possible frames and selecting sequences with stop and start codons to create spurious shadow ORFs and spurious frameshift protein sequences. The reference bacterial genomes were selected by randomly sampling 10 proteomes (Table 1) with BUSCO (Feron and Waterhouse 2022) score >95% and labelled “standard” by Complete Proteome Detector (CPD) (UniProt Consortium 2021), to select for complete and good quality proteomes. 96880 synthetic sequence were produced and were then filtered to remove any which could possibly be real proteins, by filtering any with matches to Pfam, and also filtering any with sequence matches to bacterial reference proteomes (UniProtKB 2025_03) using Diamond protein sequence search (Buchfink et al. 2015). Sequences were also clustered using MMseqs2 as before, to keep only sequences from unique clusters. 91855 AntiFam-like sequences passed these filtering steps, from which 150 per organism was sampled, normalising across length bins. The structures for the synthetic sequences were predicted using AlphaFold2, AlphaFold3 and ESMFold and the confidence scores were recorded.

**Table 1.**
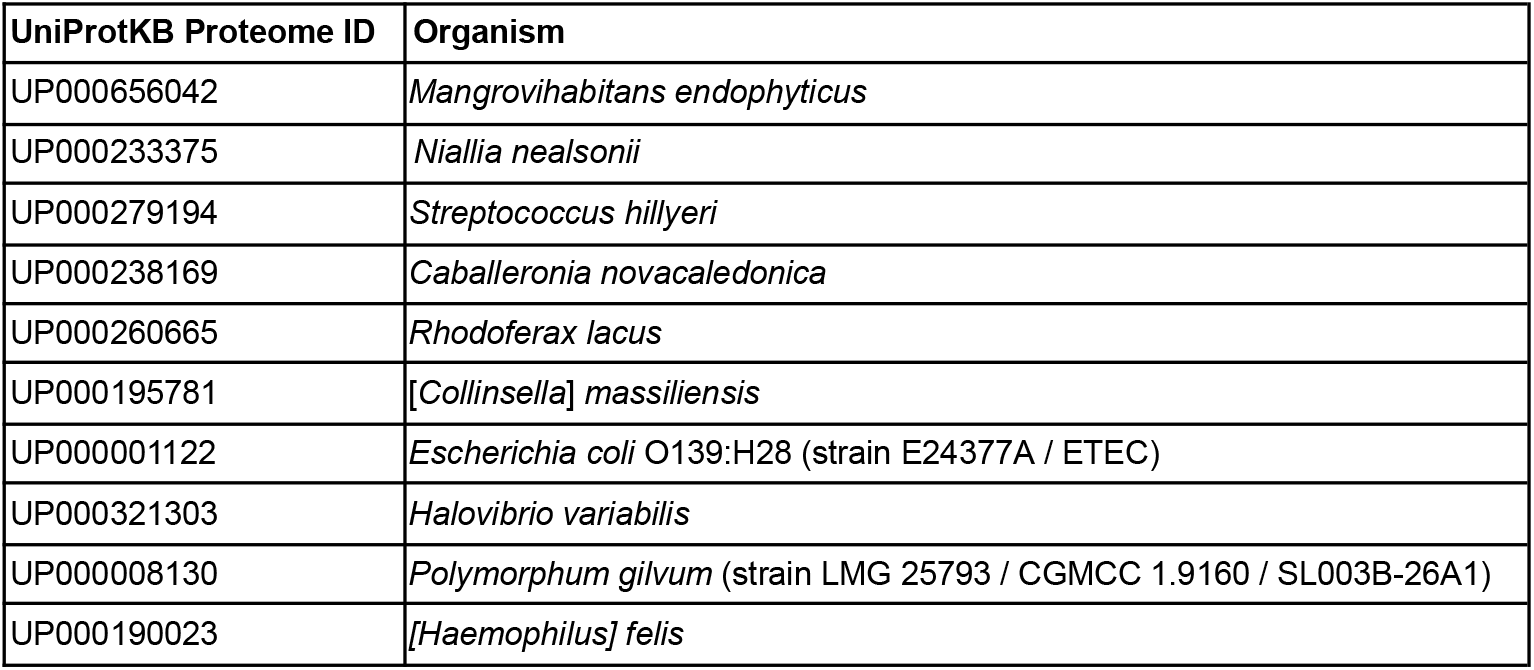
UniProtKB Proteome IDs of bacterial reference genomes sampled to generate AntiFam-like sequences.

The GPC models were tested using a balanced set with a random sample of 300 synthetic AntiFams and 300 Swiss-Prot proteins (50% spurious), as well as an unbalanced set with 100 synthetic AntiFams and 4,900 Swiss-Prot proteins, to better reflect the estimated real-world occurrence of spurious sequences (2% spurious). By default, GPC models use a threshold of 0.5. We optimised the classification threshold to maximise precision on the 2% test dataset before further application.

To apply the model to bacterial proteins with AlphaFold2 structures in AlphaFold DB, we extracted the requisite confidence scores, mean pLDDT and calculated an PAE-derived pTM from the PAE matrix of each protein structure using the following calculation:

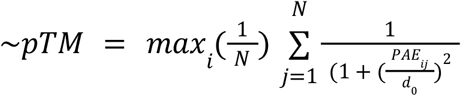

where the distance scaling factor *d*_0_ (Zhang and Skolnick 2005) is defined as in AlphaFold2 (Supplementary equation 33 in (Jumper et al. 2021)):

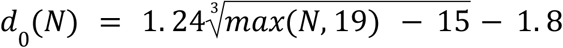

*N* is the number of residues and *PAE*_*ij*_ is the predicted aligned error between residues *i* and *j*.

PAE-derived pTM scores closely matched ColabFold pTM scores (R^2^ > 0.9; Supplementary Fig. 1). The model, designated p-AF2-GPC where p stands for PAE-derived pTM, was trained using the same dataset of AntiFam and Swiss-Prot sequences used for the previous AlphaFold2 GPC model (AF2-GPC).

Only proteins which were present in both AlphaFold DB v6 and UniProt 2025_03 were used.

## Results

### Investigation of structure prediction method confidence scores

We first examined whether ESMFold and AlphaFold3, like AlphaFold2 also produced surprisingly high pLDDT values for short spurious sequences (Fig. 1A).

**Figure 1.**
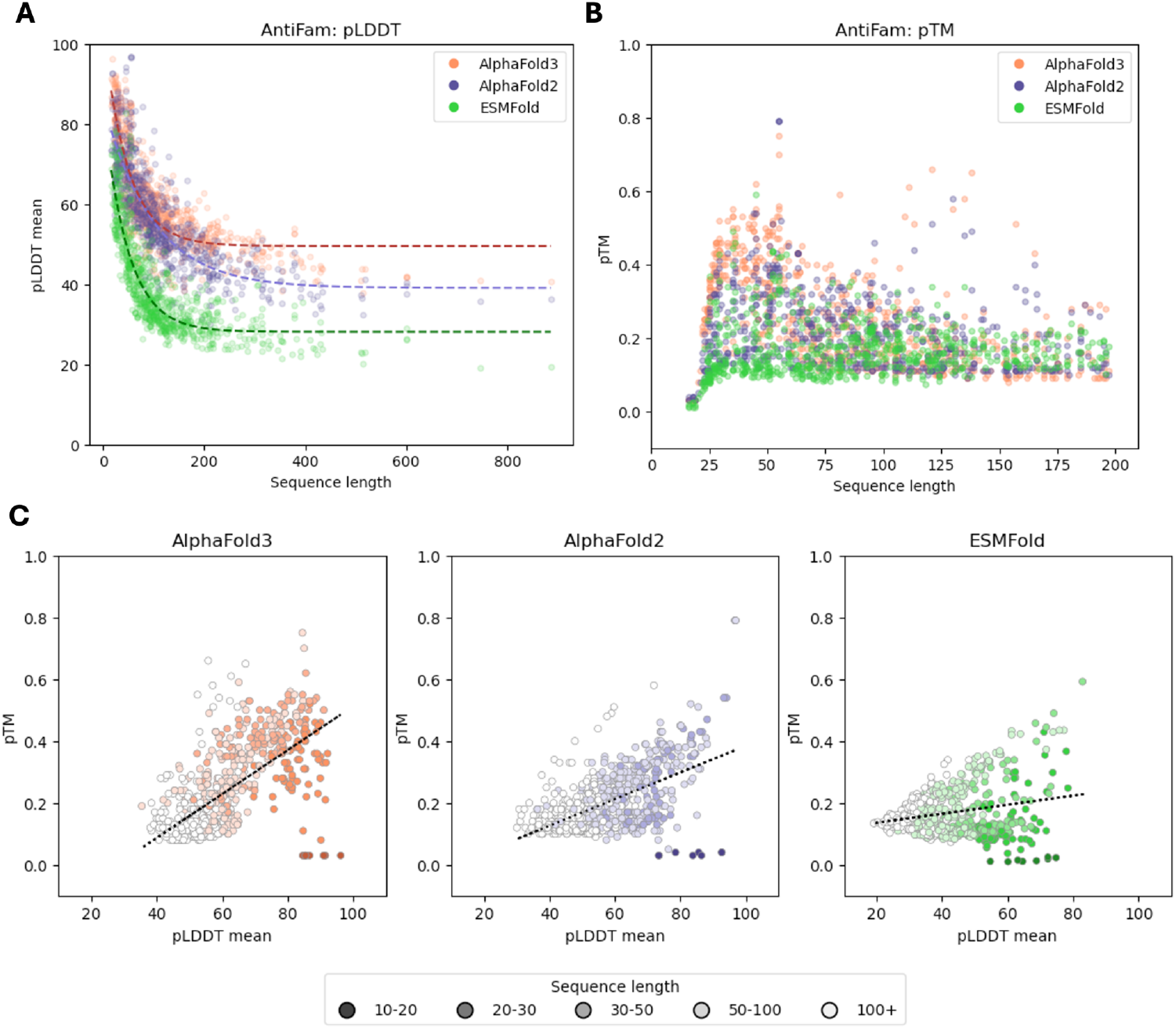
Confidence metrics for structures predicted from AntiFam sequences by ESMFold, AlphaFold2, and AlphaFold3. (A) Mean pLDDT scores by sequence length across all prediction methods. (B) pTM scores by sequence length across all prediction methods. (C) pTM versus mean pLDDT for each prediction method separately, coloured by sequence length.

All three methods predicted structures with high mean pLDDT scores above 60 for sequences shorter than 100 amino acids. AntiFam sequence lengths average at 120 and range from 16 to 886 amino acids. Of the top 10 structures with highest pLDDT scores (ranging from 91.7 to 96.8), 7 were produced by AlphaFold2 and 3 were produced by AlphaFold3. The average mean pLDDT was 60.78 for AlphaFold3, 57.36 for AlphaFold2, and 39.82 for ESMFold. The range of mean pLDDT values was most similar between AlphaFold2 (30.0 - 96.9) and AlphaFold3 (35.4 - 96.2), while ESMFold predicted lower confidence scores overall (19.1 - 82.9).

For pTM versus length (Fig. 1B), all three methods show a clustering of sequences with low pTM (under 0.05), for protein lengths below 25. AlphaFold2 and AlphaFold3 predicted some protein structures with high pTM scores (over 0.5) compared to equivalent-length sequences, as shown by the outlying points in Fig. 1B. For sequences longer than 60 residues, most pTM scores ranged from 0.1 to 0.4.

When examining the correlation between mean pLDDT and pTM, ESMFold showed the weakest linear correlation (Fig. 1C). AlphaFold3 showed the strongest correlation between mean pLDDT and pTM (0.65), while AlphaFold2 showed a moderate correlation (0.56), similar to previous observations (Monzon et al. 2022). For all three methods, it was also observed that shorter sequences (under 20 residues) had consistently low pTM (under 0.1), but high pLDDT, with AlphaFold3 predicting these short sequences with very high pLDDT, greater than 80.

Comparing AntiFam, Random and Swiss-Prot structure predictions, Swiss-Prot proteins generally were predicted with higher confidence (pLDDT or pTM) (Fig. 2), however below 100 residues, there was less separation between sequence types. Notably, AlphaFold2 and AlphaFold3 predicted Random sequences with lower confidence than AntiFam sequences, while ESMFold which does not use MSAs showed similar confidence for both sequence types.

**Figure 2.**
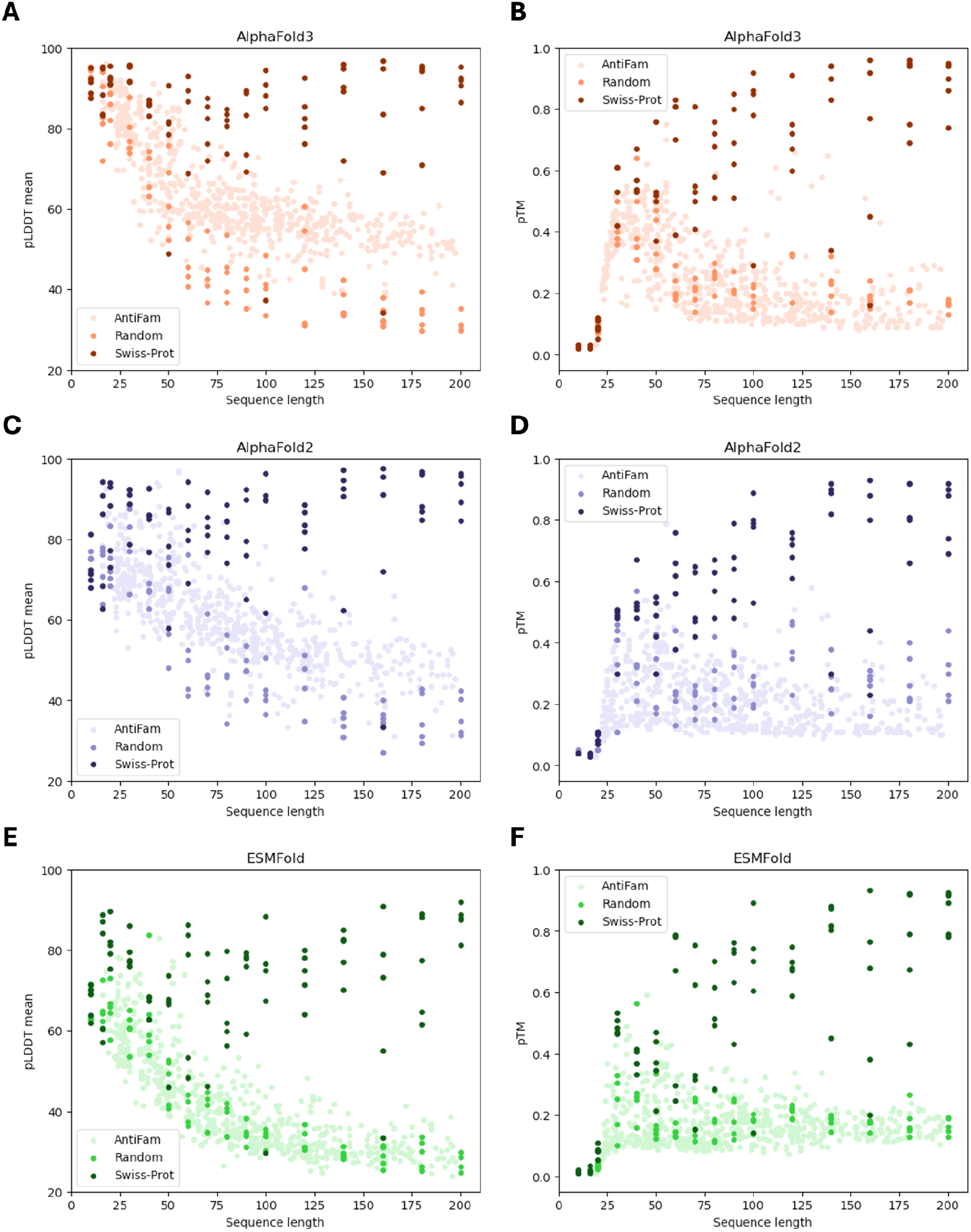
Comparison of prediction confidence across structure prediction methods, using AntiFam, Random and Swiss-Prot sequences. Mean pLDDT across sequence lengths for (A) AlphaFold3, (B) AlphaFold2, and (C) ESMFold. pTM across sequence lengths for (D) AlphaFold3, (E) AlphaFold2, and (F) ESMFold.

Interestingly, there were also several Swiss-Prot predictions with lower confidence than spurious sequences. For example, among the AlphaFold2 predictions, one long Swiss-Prot sequence (160 residues, UniProtKB: P0CX99) exhibited a pLDDT value below 40, lower than AntiFam sequences of equivalent length. This was also observed in AlphaFold3 predictions, where two additional Swiss-Prot sequences (100 residues, UniProtKB: Q12115, and 50 residues, UniProtKB: Q9GKQ1) displayed lower pLDDT values than both AntiFam and some Random sequences of comparable length.

ESMFold predicted five Swiss-Prot proteins with pLDDT below 50, (UniProtKB accessions: B8GJW5, P0CX99, P44107, Q12115, Q9GKQ1). AlphaFold3 predicted 3 proteins to have pLDDT values less than 50 (P0CX99, Q12115, Q9GKQ1), while AlphaFold3 only predicted P0CX99 with pLDDT less than 50.

P0CX99 was inferred to be a protein based on sequence similarity, and is also part of a curated family (UPF0479), however all other members in the family have only putative evidence and are from the same yeast strain, *Saccharomyces cerevisiae* S288c. A blastp search (Altschul et al. 1990) revealed no other homologs outside of *S. cerevisiae*, suggesting this protein may not be conserved in related species. P0CX99 was annotated as a membrane protein and was also found to have sequence matches completely overlapping with YRF1-4, suggesting it may be a spurious shadow ORF. Further inspection of the S288c strain reference genome (Engel et al. 2024), revealed another 18 nearly identical (>90% identity) copies of the gene, with 17 out of 19 labelled as “dubious”, many of which overlap with other ORFs. The remaining two, including P0CX99, were described as “Uncharacterised ORFs”. The protein-coding evidence for P0CX99 (locus YFL068W) was “SWAT-GFP and mCherry fusion proteins localize to the cytosol.”

Following the discovery of P0CX99 as a potential shadow ORF, we screened the rest of the low pLDDT Swiss-Prot proteins to find overlapping ORF matches in other reading frames. We found Q12115, also from *Saccharomyces cerevisiae S288c*, to be a shadow ORF in the -2 frame of CDC7 (P06243), with half of the Q12115 ORF overlapping at the C-terminus. Q12115 is also marked by UniProtKB as a dubious gene prediction.

### Identification of a potential false-positive AntiFam

We inspected the structures of AntiFam structures with the highest pTM and pLDDT for globular, protein-like structures. The AntiFam ANF00264 predictions had the highest pTM and were also one of the AntiFam predictions with high pLDDT which were not a single alpha-helix. We tested whether this protein might form a homodimer by generating a dimeric structure with AlphaFold2 (Fig. 3B). The PAE plot supported interactions between the two homodimeric subunits (Fig. 3C). The mean pLDDT was 94.8, the pTM was 0.881, and the ipTM was 0.87. No other AntiFam entries were found to have globular structure.

**Figure 3.**
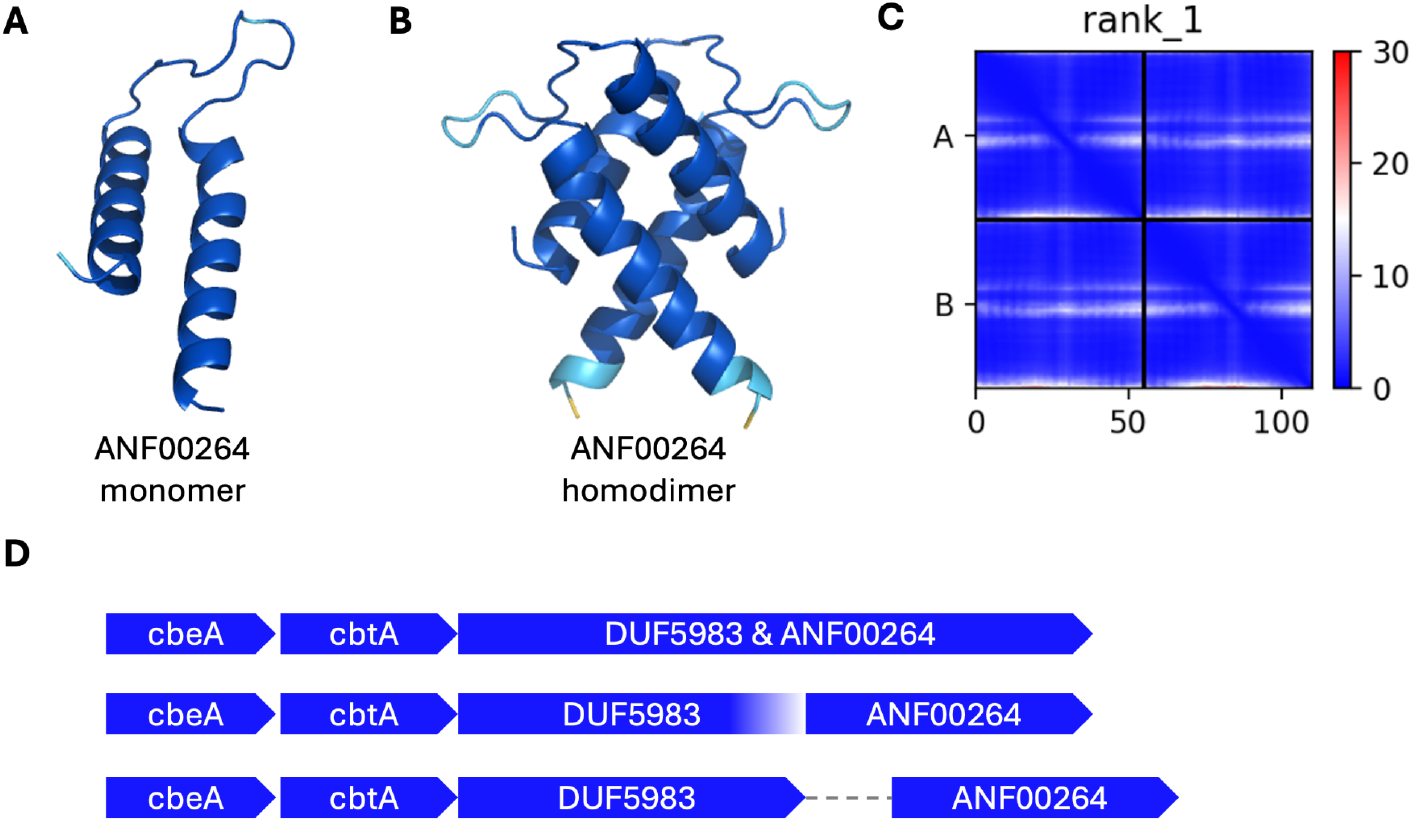
AlphaFold2 structure predictions of potential non-spurious AntiFam entry ANF00264. ANF00264 monomer, coloured by pLDDT (A). ANF00264 homodimer, also coloured by pLDDT (B). Predicted Aligned Error (PAE) plot of ANF00264 homodimer to show confidence of relative position of residues and domains within the structure (C). Local genomic contexts of ANF00264: DUF5983 and ANF00264 predicted as part of a single protein, incomplete C-terminus DUF5983 and ANF00264 predicted as separate proteins, and distinct DUF5983 and ANF00264 predicted proteins separated by a non-coding region (D).

AntiFam describes ANF00264 as a set of spurious proteins derived from frameshifted phage proteins. Although frameshifts often indicate spuriousness, programmed frameshifts have been documented in several Bacteriophage proteins (Sipley et al. 1991), (García et al. 2004), (Xu et al. 2004), raising the possibility that ANF00264 could represent a genuine programmed frameshift protein.

A blastp search performed using the first seed sequence of ANF00264 returned a 100% match to a hypothetical protein (UniProtKB: A0A083ZZA7) found in Serratia Sp., as well as more distant matches in other Enterobacteriaceae species. We examined the domains of this protein and found it had matches to ANF00264 as well as the Pfam domain, “Family of unknown function (DUF5983) C-terminal domain” (Pfam: PF19419), which is also described as likely phage derived (Paysan-Lafosse et al. 2025). Extending this search, we also identified a further 24 proteins containing both ANF00264 and DUF5983 matches. Among these proteins, 8 are named as CP4-44 prophage proteins and 4 as Aec77 proteins. These proteins occur in multiple bacterial species including *Escherichia coli, Klebsiella pneumoniae*, and *Shigella flexneri*. Notably one of these proteins (UniProtKB: A0A447 × 825) also contained the CbtA toxin domain (Pfam: PF06755).

CbtA (Cytoskeleton binding toxin A) inhibits cell division by binding to cytoskeleton proteins FtsZ and MreB, ultimately causing cell death. CbtA functions with the antitoxin CbeA within a toxin-antitoxin (TA) system, where both genes are typically encoded adjacently within the same operon (Brown and Shaw 2003).

This finding of ANF00264 adjacent to a well-characterised operon, prompted us to examine whether this was an isolated case of a conserved feature in other bacteria with the CbtA system. We found that the proteins containing the ANF00264 and DUF5983 matches were consistently found to be encoded next to cbtA and cbeA genes.

Interestingly, ANF00264 did not always appear as part of a multidomain protein with a DUF5983 domain. Sometimes it was predicted as a separate protein, sometimes separated by a short non-coding region. We hypothesise that this difference reflects the programmed frameshift, where annotation tools may predict a fused or separate protein. This is also possible given assemblies where DUF5983 protein upstream of ANF00264 was annotated to have an incomplete C-terminus, resulting in two distinct predicted proteins.

The repeated association of ANF00264 with a functional operon, despite variation in annotation, suggests it is not a spurious sequence but instead may be a phage-derived domain or protein associated with the CbtA TA system.

### Evaluation of Gaussian Process Classification (GPC) models trained with structure prediction results

To test whether prediction confidence scores could be used to systematically distinguish between real and spurious proteins, we trained three GPC models (AlphaFold2-GPC (AF2-GPC), AlphaFold3-GPC (AF3-GPC) and ESMFold-GPC (ESM-GPC). The models used three features, sequence length, and confidence scores mean pLDDT and pTM which were derived from each respective structure prediction method.

To evaluate the classification performance of the model, we carried out receiver operating characteristic (ROC) analysis to assess the classifiers’ performance in identifying spurious protein sequences. The ROC curves were generated by plotting the true positive rate (recall) against the false positive rate (Fig. 4A). This approach helps in visualising the trade-off between correctly identifying spurious sequences (true positives) and incorrectly classifying real sequences as spurious (false positives). The area under the ROC curve (AUC) summarises the ROC into one value, representing the probability that the classifier will score a random spurious sequence higher than a random real sequence.

**Figure 4.**
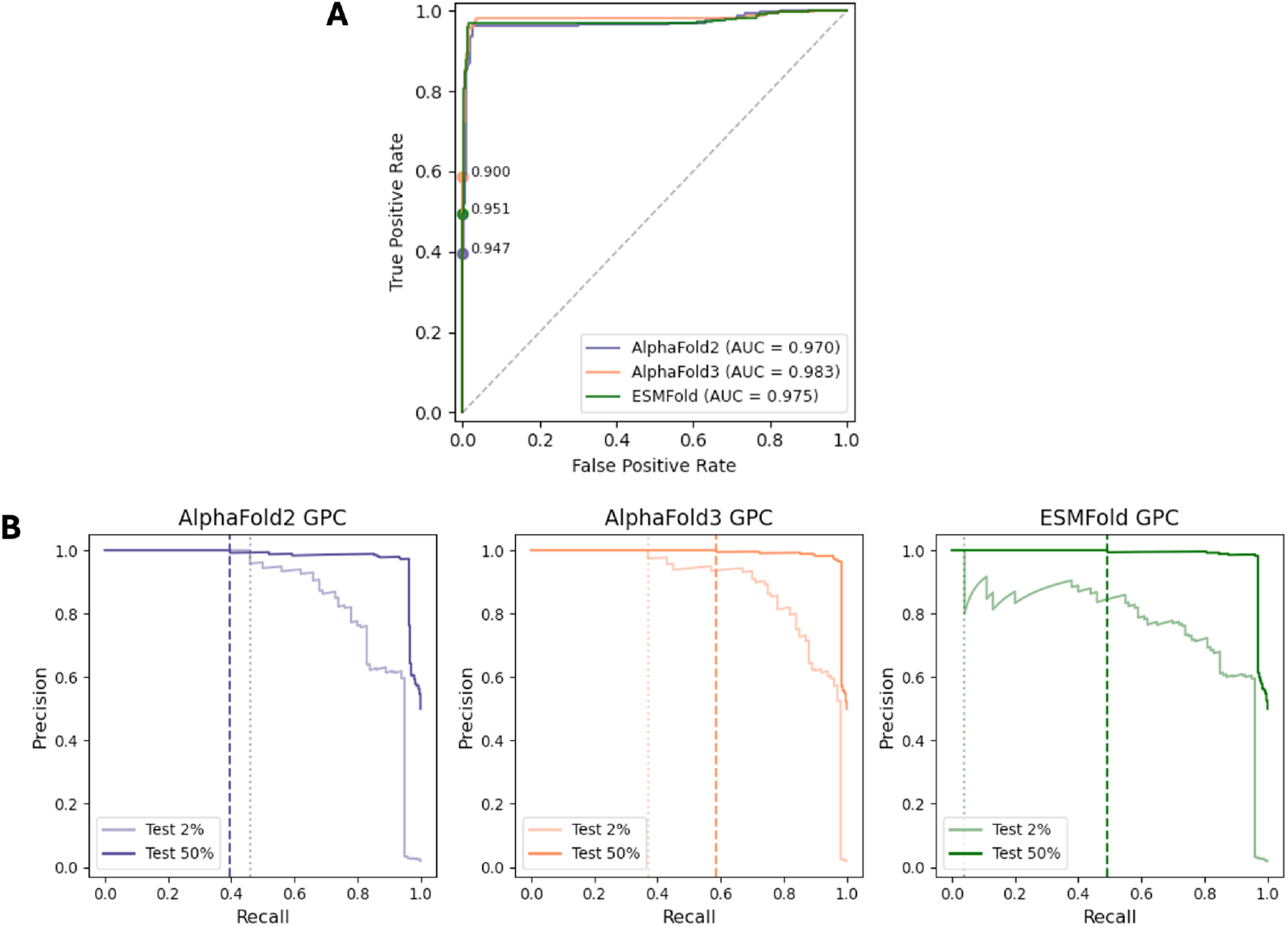
(A) ROC curves for GPC models testing on balanced datasets (50% synthetic AntiFam structure prediction scores, 50% Swiss-Prot structure prediction scores). (B) Precision-recall curves for GPC models. Models were tested on balanced datasets constructed from 50% synthetic AntiFam structure predictions and 50% Swiss-Prot structure predictions, and unbalanced datasets with 2% synthetic AntiFam structure predictions and Swiss-Prot structure predictions, shown by the darker and lighter lines. Dashed lines mark maximum recall at 100% precision using the unbalanced test set, and dotted lines mark maximum recall at 100% precision using the balanced test set.

**Figure 5.**
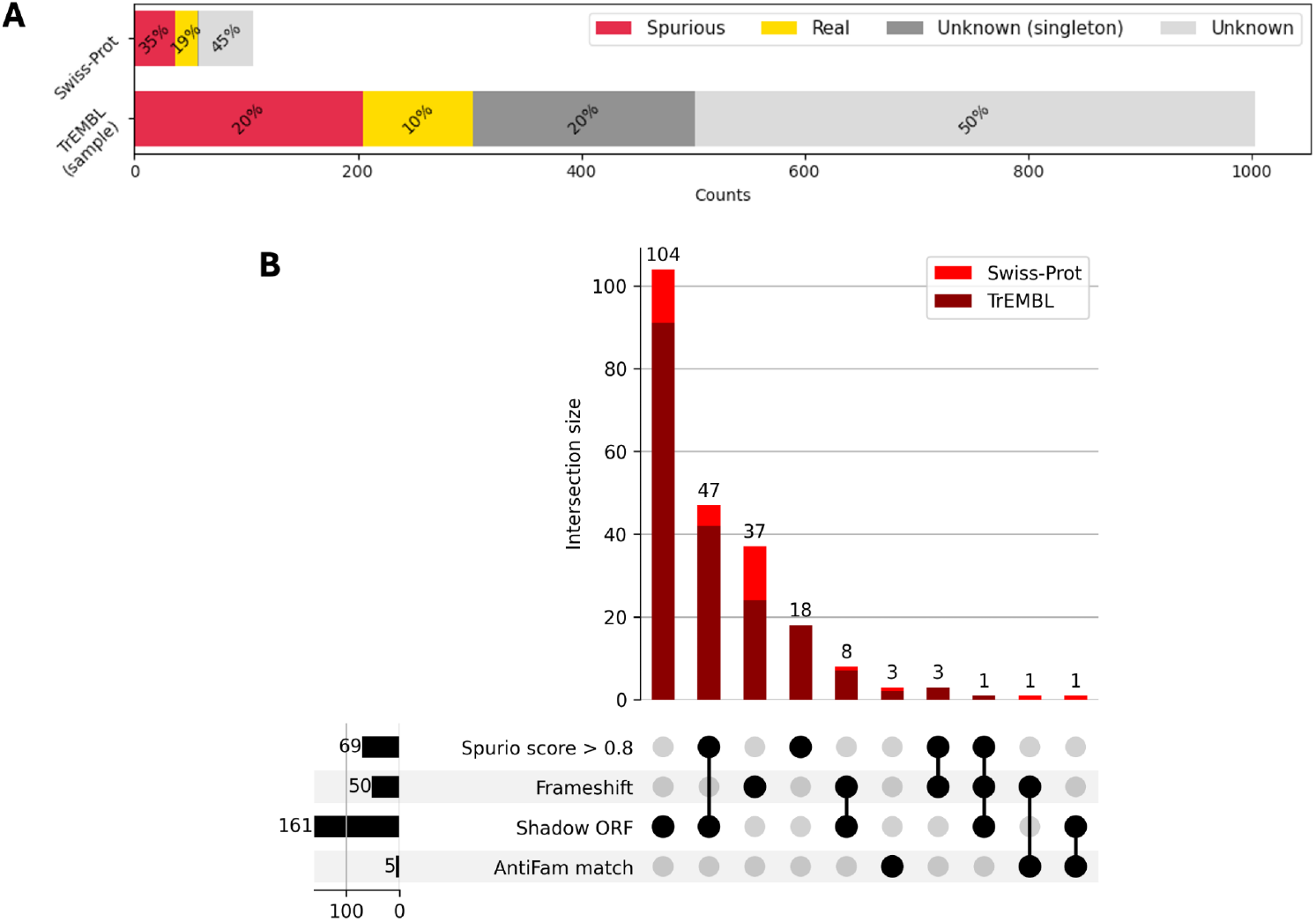
(A) Investigation results for proteins predicted as spurious by p-AF2-GPC: all spurious Swiss-Prot proteins and a random sample (1000) of predicted spurious TrEMBL proteins. Additional evidence was used to assess the proteins as spurious, real proteins (false positives), and unknown (insufficient evidence to confirm or reject prediction). Unknown proteins were subdivided into singleton proteins (potential orphan genes with few or no sequence matches). (B) UpSet plot showing the overlap of four additional spurious protein detection methods used to validate p-AF2-GPC predictions. Horizontal bars show total number proteins predicted as spurious by each method. Vertical bars show the number of proteins in each intersection and the stacks are coloured to distinguish Swiss-Prot (red) and TrEMBL (dark red).

Model precision was prioritised over recall, to minimise incorrectly classing real proteins as spurious. We generated precision-recall curves for each model (Fig. 4B) to identify if the models maintained reasonable recall rates, if optimised for 100% precision.

We evaluated the models using both a balanced test set and an unbalanced dataset, reflecting the occurrence of spurious sequences in protein databases (Fig. 4B). When tested on the balanced dataset (50% spurious), AF3-GPC demonstrated the best performance with 59% recall at 100% precision. ESM-GPC achieved 49% recall at 100% precision, followed by AF2-GPC with 40% recall at 100% precision. When tested on the unbalanced dataset (2% spurious), both AF3-GPC and ESM-GPC models experienced a decrease in recall at 100% precision. The AF3-GPC recall at 100% precision decreased to 37%. The ESM-GPC recall at 100% precision decreased greatly to 4%. Interestingly, the AF2-GPC performance improved slightly when tested on the unbalanced set with 46% recall at 100% precision.

### Applying classification model on AlphaFold DB

We also explored the possibility of applying the AF2-GPC on AlphaFold DB (Varadi et al. 2024), a large-scale database of structure predictions. As AlphaFold DB lacks pTM scores, we calculated PAE-derived pTM scores from the available PAE matrices (see Methods). Using PAE-derived pTM, mean pLDDT and length as features, we trained a model designated p-AF2-GPC on the same AntiFam and Swiss-Prot datasets used for AF2-GPC. When tested on the unbalanced test set, p-AF2-GPC achieved 32% recall at 100% precision, with a classification threshold of 0.957.

AlphaFold DB consists of sequences from two main sources: UniProtKB reviewed entries (Swiss-Prot), which represent manually curated, high-quality protein sequences and UniProtKB unreviewed entries (TrEMBL), which are computationally annotated and may contain errors or spurious sequences. We focused on bacterial proteins for two main reasons; our collection of spurious families (AntiFam) are largely from bacterial sequences, and eukaryote proteins more commonly exhibit disordered regions (Basile et al. 2019) which reduces structure prediction confidence, which may lead to false positive predictions. AlphaFold DB has 115,057,521 bacterial TrEMBL protein structure predictions, and 335,771 bacterial Swiss-Prot protein structure predictions.

Applying p-AF2-GPC on bacteria-only-AlphaFold DB returned 378,358 TrEMBL proteins (0.3% of bacterial TrEMBL proteins) and 105 Swiss-Prot proteins (0.03% of bacterial Swiss-Prot proteins) predicted as spurious. While the number of Swiss-Prot proteins is low, further investigation was performed to investigate if all 105 were indeed spurious proteins.

The proteins predicted as spurious were assessed to be likely real or spurious with the following criteria: proteins were classified as real if they have Pfam matches, or experimental evidence at the protein level. Protein evidence scores were obtained from the UniProtKB entry annotations for each protein. The scores scale from to 1-5 depending on strength of existence evidence where 1 is direct protein detection. Proteins were classified as spurious if they have AntiFam matches, score highly with tools such as Spurio (Höps et al. 2018), or are found to be frame mispredictions of well-known proteins (shadow ORF or frameshift sequences).

Out of the 105 Swiss-Prot proteins p-AF2-GPC predicted as spurious, 20 were found to have at least one match to a Pfam family and or experimental evidence, and therefore were likely real proteins. A quarter of these were also classed as disordered by MobiDB consensus (Piovesan et al. 2025). 37 of the Swiss-Prot proteins were assessed to be likely spurious as they either had AntiFam matches, Spurio score >0.8, shadow ORF or frameshift evidence.

The TrEMBL proteins are harder to classify, as the large majority are not experimentally characterised. However much of TrEMBL is still covered by Pfam, 76% of proteins have a match to Pfam. We randomly sampled 1,000 of the TrEMBL proteins predicted by the model to be spurious to gauge if there was a similar false positive prediction rate. From the sample we found 98 proteins with Pfam matches, and out of these real proteins, 36 were predicted to be intrinsically disordered. This left 700 unclassified proteins, 405 of which were predicted to be intrinsically disordered. 205 proteins from the TrEMBL sample were assessed to be likely spurious, either having AntiFam matches, Spurio score >0.8, shadow ORF or frameshift evidence.

We classed proteins to be likely spurious if they could also be validated using an additional spurious protein detection method, such as Spurio, AntiFam or by finding sequence matches in alternative reading frames (shadow ORF or frameshift evidence). The largest group consisted of 104 proteins with shadow ORF evidence and the second largest group consisted of 47 proteins with both shadow ORF evidence and high Spurio score (>0.8), representing high-confidence spurious predictions. These 47 proteins were compiled along with other candidates with high Spurio score and frameshift evidence into a dataset of confident spurious predictions (https://github.com/0rra/fold_unfold2/blob/main/part3_gpc_testing/3_results_plotting/results/spurious_candidates.tsv). There were few AntiFam matches (5 proteins in total, including 3 from Swiss-Prot), suggesting most of the validated spurious proteins are novel findings not yet represented in AntiFam.

## Discussion

We found that ESMFold, AlphaFold2 and AlphaFold3 all predict short sequences with high pLDDT values, regardless of whether the sequences represent spurious or randomly generated sequences. As a result, significant separation in pLDDT between spurious and real sequences only becomes apparent when sequence length exceeds 100 amino acids.

A likely cause for the length bias could be there are fewer possible structural arrangements for shorter sequences to adopt, therefore can more easily be predicted, contributing to higher pLDDT values even when the underlying sequence may be spurious. Very short sequences under 30 amino acids, real and spurious, are also predicted with very low pTM scores, as well as high pLDDT. To further investigate this pattern in confidence scores for short sequences and the seemingly limited structural variation (e.g. alpha-helix structure), we can inspect the predicted structures of a larger pool of experimentally verified short proteins.

Out of the 80 randomly sampled proteins from UniProtKB/Swiss-Prot, two were found to be likely spurious proteins. Both shadow ORF proteins were over 100 amino acids long (100 and 160 in particular), consistent with our finding that structure prediction confidence becomes more discriminative above this threshold.

We also observed that AlphaFold2 and AlphaFold3 differentiated Random sequences from AntiFam sequences, predicting Random sequences with lower confidence. This likely stems from their architectural design, as both methods use multiple sequence alignment (MSA), and the lower number of matches in the MSA for random sequences results in lower confidence predictions. AntiFam sequences may have higher coverage than random sequences due to matching with other spurious protein sequences remaining in sequence databases. ESMFold does not use an MSA, perhaps explaining why it does not differentiate between AntiFam and Random sequences.

While the GPC model has a non-zero false positive prediction rate, it can effectively enrich candidate spurious proteins, combined with additional methods such as Spurio. The sampled flagged TrEMBL proteins included 20% spurious sequences, compared to the predicted 1-2% in TrEMBL overall. While the model does misidentify some real proteins, only 10% of the flagged TrEMBL proteins had Pfam matches versus the 76% Pfam coverage of TrEMBL overall. This suggests the model successfully enriches spurious proteins in its output. The model could be used to quickly narrow down potential spurious proteins given structure prediction confidence scores, and be used in combination with more time-consuming methods to further validate and identify spurious proteins with greater confidence.

The remaining unexplained proteins without Pfam matches may represent genuine but uncharacterised proteins, sequences with few to no homologs (orphan genes) or spurious sequences which evade current detection methods. Limited homology may indicate either novelty *i*.*e. de novo* gene birth, or lack of evolutionary conservation as would be expected of spurious sequences. A similar issue remains with intrinsic disorder, as this property appears coincidentally with spurious protein sequences.

In conclusion, we have shown that all structure prediction methods tested predict short sequences with high pLDDT. This hampers our ability to use structure prediction metrics to identify spurious proteins which are often short. We present a Gaussian Process Classifier method which distinguishes real and spurious proteins imperfectly, but it is fast to apply and can be used as a pre-filtering step before using more computationally expensive methods to identify spurious proteins. The identification of spurious proteins remains an important but unsolved challenge in protein bioinformatics.

## Funding

A.K.O and A.B. were supported by core EMBL funding.

## Software and Data availability

The structure predictions for all sequences are available at https://doi.org/10.5281/zenodo.18390113, and the structure prediction NextFlow pipeline and code used for figure generation can be found at https://github.com/0rra/fold_unfold2.

## Notes

### Competing Interest Statement

The authors have declared no competing interest.

