## Supplementary figures and images for "Folding the unfoldable 2: using AlphaFold and ESMFold to explore spurious proteins"

### Supplementary Fig. 1

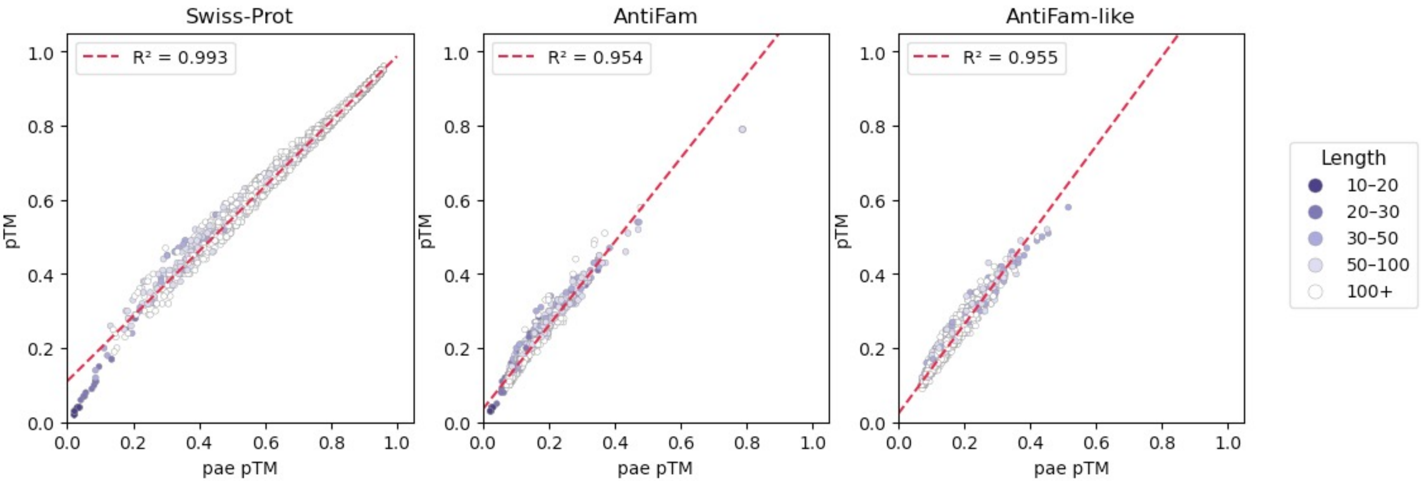
